# The Fragile X Mental Retardation Protein protects the lung from xenobiotic stress by facilitating the Integrated Stress Response

**DOI:** 10.1101/2021.03.16.435743

**Authors:** Deblina Sain Basu, Rital Bhavsar, Imtiyaz Gulami, Sai Manoz Lingamallu, Ravi Muddashetty, Chandrakanth Veeranna, Sumantra Chattarji, Rajesh Thimmulappa, Aditi Bhattacharya, Arjun Guha

## Abstract

Stress response pathways protect the lung from the damaging effects of environmental toxicants. Here we investigate the role of the Fragile X Mental Retardation Protein (FMRP), a multifunctional protein implicated in stress responses, in the lung. We report that FMRP is expressed in murine and human lungs, in the airways and more broadly. Analysis of airway stress responses in mice and in a murine cell line *ex vivo,* using the well-established Naphthalene (Nap) injury model, reveals that FMRP-deficient cells exhibit increased expression of markers of oxidative and genotoxic stress and increased cell death. We find that FMRP-deficient cells fail to actuate the Integrated Stress Response Pathway (ISR) and upregulate the transcription factor ATF4. Knockdown of ATF4 expression phenocopies the loss of FMRP. We extend our analysis of the role of FMRP to human bronchial BEAS-2B cells, using a 9, 10-Phenanthrenequinone air pollutant model, to find that FMRP-deficient BEAS-2B also fail to actuate the ISR and exhibit greater susceptibility. Taken together, our data suggest that FMRP has a conserved role in protecting the airways by facilitating the ISR.

## INTRODUCTION

The epithelial lining of the respiratory tract is continually challenged by a diverse array of environmental toxicants including gases, particulates, and other biological agents. Exposure to these agents leads to increased oxidative, genotoxic and endoplasmic reticulum stress. Such stresses, when unmitigated, lead to cellular damage, inflammation and in the long term to decreased lung capacity and functionality. The aim of this study was to probe the mechanisms by which lungs cope with environmental stresses.

The capacity of the lung to manage xenobiotic stress is dependent on stress response proteins that are induced upon insult. In this regard, the Integrated Stress Response (ISR) pathway is an evolutionarily conserved pathway that is integral to how the lung copes with environmental challenges (Pakos-Zebrucka K et al., 2016; van ’t Wout EF et al., 2014; Konsavage WM et al., 2012). The ISR is triggered by the activation of one or more of the four stress-responsive kinases GCN2, PKR, PERK and HRI. The activation of these kinases, in turn, sets in motion two separate but interdependent processes that enable cells to mount a restorative response (Wong HR & Wispe JR, 1997). First, these kinases phosphorylate the Eukaryotic Initiation Factor 2α (eIF2α) and shut off ongoing programs of protein synthesis. The inhibition of translation leads to the sequestration of translationally active mRNAs into stress granules (SGs). Second, activation of the kinases also induces specialized modes of protein translation leading to the expression of stress response proteins. More specifically, these specialized translation regimes upregulate expression of Activating Transcription Factor 4 (ATF4) (Pakos-Zebrucka K et al., 2016; van ’t Wout EF et al., 2014) and, in turn, ATF4 targets such as ATF3. ATF4 also synergizes with other transcription factors activated in response to stress like Nrf2, to induce the expression of other stress response genes (He CH et al., 2001; Sarcinelli C et al., 2020).

The Fragile X Mental Retardation Protein (FMRP) is a multifunctional protein that is expressed in the brain and more widely, in humans and other mammals alike. Deficiencies in FMRP lead to Fragile X Mental Retardation Syndrome (FXS), a disease characterized by mild-to-moderate intellectual disability (Zhou Z et al., 2014). FMRP function has been most intensively studied in the neuronal context wherein the protein has been shown to regulate synaptic plasticity by multiple mechanisms (Santoro MR et al., 2012). Aside from this well-established role, several studies indicate that FMRP also has a role in facilitating stress responses. At a cellular level, FMRP has been shown to play an essential role in SG biogenesis in response to arsenite and heat shock-induced stress (Didiot MC et al., 2009; Linder B et al., 2008). A recent study on fibroblasts in FMR1 KO mice showed that these cells are unable to mount a DNA Damage Response (DDR) when exposed to agents like Aphidicolin, 5-Hydroxyurea (5-HU) and UV but are able to do so in response to other types of DNA damaging agents (Alpatov R et al., 2014). The central finding of this study is that FMRP has a chromatin-dependent role in resolving stalled replication forks and single strand breaks in DNA (Alpatov R et al., 2014). The environmental toxicants that the lung is exposed to typically induce a wide spectrum of genotoxic perturbations. Whether the chromatin-dependent role of FMRP is essential in this milieu is not clear.

Our interest in candidate proteins that regulate the pulmonary stress response led us to explore the role of FMRP in the lung. Immunostaining of murine and human lungs revealed that the protein is expressed in the airway epithelium and more broadly. To probe the role of FMRP in stress responses in the airways, we subjected Fmr1 KO mice to Naphthalene injury, a well-established model for oxidative and genotoxic stress. We found that the airways of Fmr1 KO mice exhibited higher expression of markers of oxidative and genotoxic stress, and greater cell death, than wild type. These findings led us to investigate the role of FMRP in airway stress responses in mice, the involvement of the protein in the ISR, and its role in the human lung.

## RESULTS

### FMRP is expressed in the airways and more broadly in the murine lung and protects airway Club cells from Naphthalene induced stress

To characterize the role of FMRP in the pulmonary stress response, we examined the expression of the protein in adult lungs from wild-type (WT) and Fmr1 KO animals. Lung sections from WT mice were stained with anti-FMRP antisera and examined under a confocal microscope (5 um, n>3 mice). FMRP expression was detected throughout the lung (Fig. 1A, 1C). We detected widespread protein expression in airway epithelium, both in secretory Club cells (CCs, marked by expression of Scgb1a1, Fig. 1A, 1C) and in ciliated cells (marked by expression of Acetylated Tubulin, AcTub). Outside of the airways, we noted intermittent expression in the alveolar parenchyma (Fig. 1A). Lung sections of Fmr1 KO mice stained with the same anti-FMRP antisera did not show any specific staining (airways shown in Fig. 1B, 1D, n>3 mice). Together, these experiments showed that FMRP is expressed in the murine lung, in the airways and more broadly. We also examined H&E stained lung sections from WT and Fmr1 KO mice to find no obvious abnormalities in Fmr1 KO lungs (Fig. S1 A-B).

**Figure 1:**
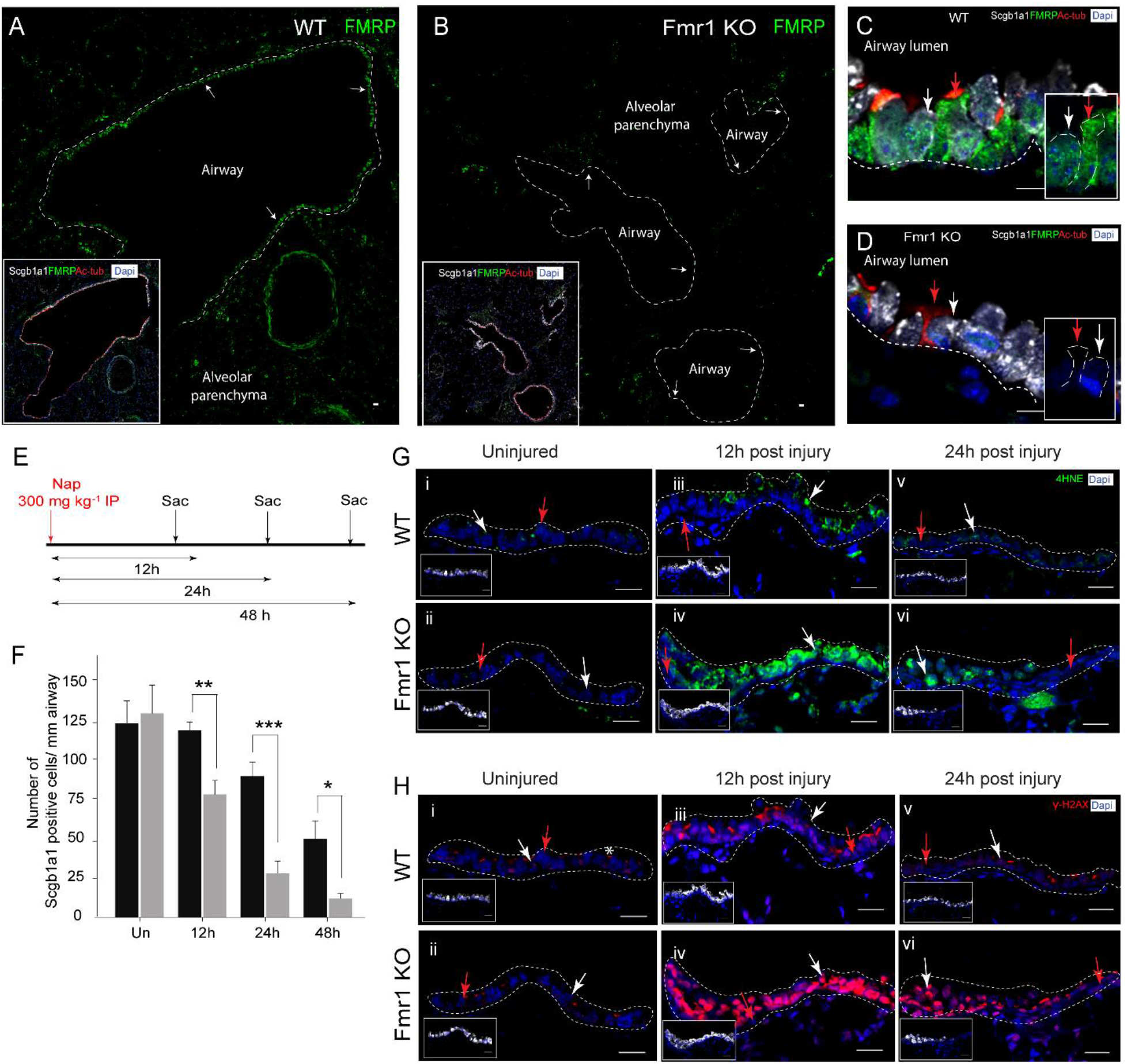
FMRP is expressed in the airways and more broadly and protects airway Club cells from Naphthalene induced stress. **(A-D)** FMRP expression in the murine lung. **(A)** Tiled image showing FMRP immunostaining (green, arrow) in the airway epithelium (demarcated by white dotted lines) and in the parenchyma of the murine lung. The airways are identified by expression of the Club cell (CC) marker Scgb1a1 (white, inset) and of the ciliated cell marker acetylated-Tubulin (red, inset). **(B)** Tiled image showing FMRP immunostaining in Fmr1 knockout (Fmr1 KO) mice. Note absence of FMRP (green) in both airway (demarcated by white dotted lines, inset) and parenchyma. **(C-D)** High resolution image of FMRP immunostaining (green) in airway epithelial cells. Shown here are CCs (white, white arrow in panel alongside) and ciliated cells (red, red arrow in panel alongside) in wild type **(C)** and Fmr1 KO **(D)**. **(E-H)** Susceptibility of CCs to Naphthalene (Nap) injury in control and Fmr1 KO. **(E)** Schematic showing regimen for Nap injury. **(F)** Frequencies of Scgb1a1^+^ cells in wild type (black bars) and Fmr1 KO (grey bars) from uninjured (Un) and Nap-injured mice at different timepoints post injury. Data represents average + SD. **(G-H)** Expression of markers of oxidative (4HNE) and genotoxic (γ-H2AX) stress in airways from wild-type and Fmr1 KO mice prior to and post Nap. (**G i-G vi**) 4HNE immunostaining (green) in the airways of wild-type (upper panel) and Fmr1 KO (lower panel) mice prior to and post Nap. **(H i-H vi)** γ-H2AX immunostaining (red) in the airways of control (upper panel) and Fmr1 KO (lower panel) prior to and post Nap. Also see Fig S1. Statistical significance was assessed by an unpaired two-tailed t-test (see methods, p< .05*, p< .01**, p< .001***). The changes in the two groups over time, across genotype and interaction parameters were also assessed by two-way ANOVA and found to be statistically significant. For Shapiro-Wilk normality test and two-way ANOVA see Table S1. Scale Bar=20 um.

To investigate the role of FMRP in the pulmonary stress response, we focused our attention on FMRP-expressing airway CCs and exposed WT and Fmr1 KO mice to a chemical that targets CCs. Airway CCs are highly sensitive to the polycyclic hydrocarbon Naphthalene (Nap) (Stripp BR et al., 1995; Van Winkle LS et al., 1995). Nap administration leads to the loss of the vast majority of CCs from the airway epithelium within 24-48 h and is a well-established model for lung injury (Guha A et al., 2014; Guha A et al., 2017). The susceptibility of airway CCs to Nap is due to the expression, in CCs, of the cytochrome P450 enzyme Cyp2f2 (Buckpitt A et al., 2002). Cyp2f2 converts Nap to Naphthalene oxide that causes DNA damage. Naphthalene oxide is also converted to Naphthoquinones that cause oxidative stress (Buckpitt A et al., 2002). The Cyp2f2 isoform that converts Nap to cytotoxic derivatives is not expressed in humans and consequently Nap does not affect humans in the same way.

We exposed WT and Fmr1 KO animals to Nap and harvested lungs for analysis at different timepoints post injury (regimen shown schematically in Fig. 1E). To assess the extent of injury, we quantified frequencies of CCs across timepoints and examined expression of markers of oxidative and genotoxic stress. We found that the frequencies of CCs in WT were significantly higher than in Fmr1 KO at 24 h and 48 h respectively (Fig. 1F, n=3 mice per genotype per timepoint). In other words, cell loss was greater in Fmr1 KOs. Next, we stained sections from mouse lung both prior to and post Nap injury with two antisera: anti-4-Hydroxynonenal (4HNE, a product of lipid peroxidation and a marker of oxidative stress) and anti-γ-H2AX (a phosphorylated histone variant that is a marker of double stranded DNA breaks and genotoxic stress). We did not detect expression of either stress marker in the lung in uninjured WT and Fmr1 KO mice (Fig. S1C, S1D, also Fig. 1G i-1G ii, 1H i - 1H ii) and the expression of both markers was dramatically increased in Nap-injured lungs. Pertinently, we noted that the frequencies of CCs were higher and the levels of 4HNE and γ-H2AX expression were lower in WT than in Fmr1 KOs, at all timepoints examined (Fig. 1G iii-1G vi,1H iii-1H vi, Fig. S1, n=3 mice per genotype per timepoint). Based on these data we concluded that CCs in Fmr1 KO animals are more susceptible to Nap-induced stress.

### The Club cell-like C22 cell line deficient in FMRP is also more susceptible to Nap induced stress

To further probe the role of FMRP in stress responses in CCs, we turned to the murine Club cell-like cell line, C22. C22 cells were isolated from H-2Kb-tsA58 mice expressing a temperature sensitive isoform of the SV40 Large T antigen under the H-2Kb promoter (Demello DE et al., 2002). To characterize these cells, we stained C22 cells with markers of CCs and other airway and alveolar lineages. Consistent with results from previous reports, these cells expressed the CC marker Scgb1a1 and did not express markers of other lineages (Fig. 2A, data not shown). To determine whether C22 could be utilized as a model for Nap injury, and to probe the role of FMRP therein, cells were stained with antisera against Cyp2f2 and FMRP. We found that C22 cells expressed modest levels of Cyp2f2 (Fig. 2B) and also expressed FMRP (Fig. 2C, n>6 experiments).

**Figure 2:**
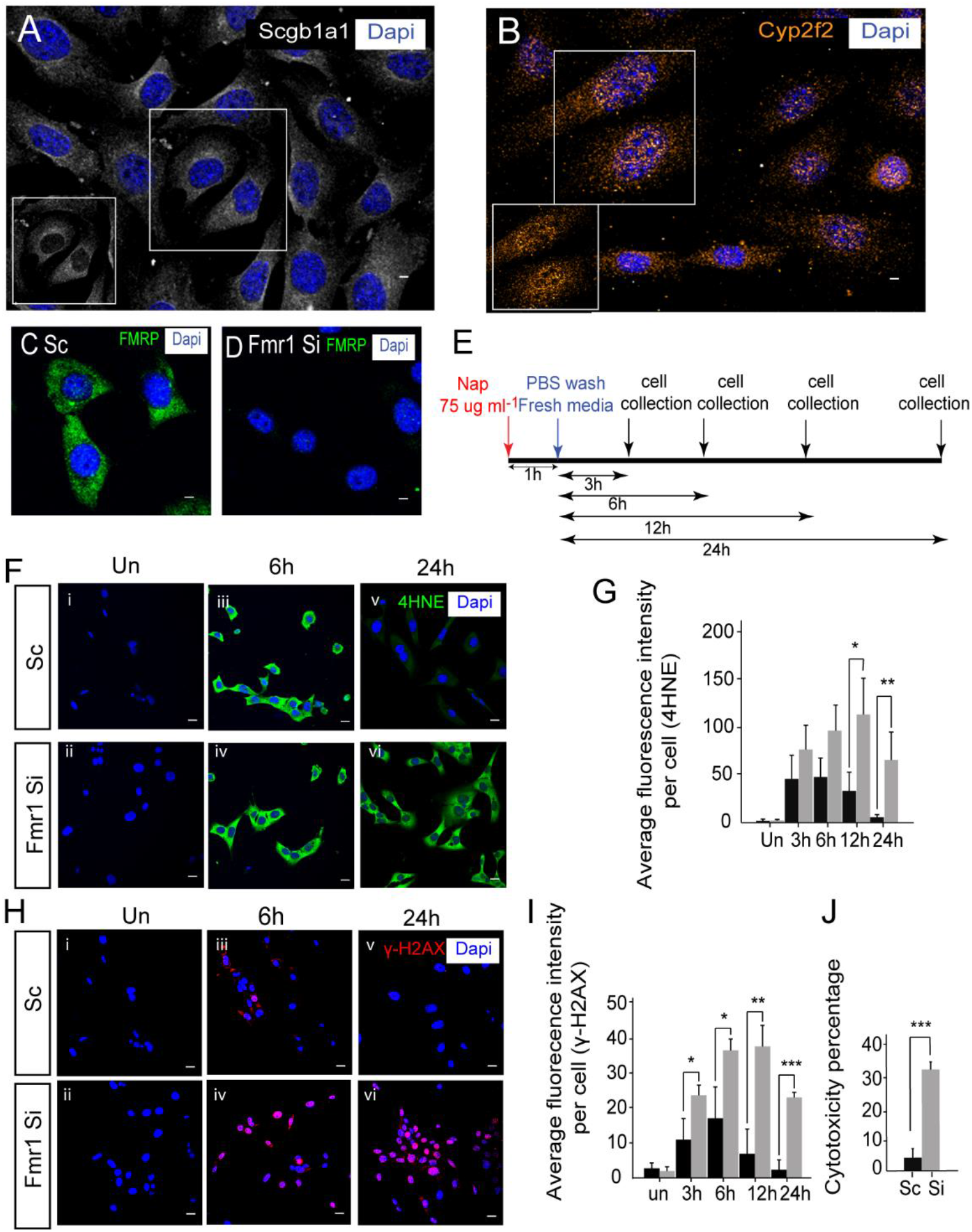
FMRP deficient Club cell-like C22 cells are susceptible to Nap-induced stress. **(A-D)** Phenotypic characterization of C22 cells. **(A)** Scgb1a1 immunostaining (white) in C22 cells. **(B)** Cyp2f2 (orange) immunostaining in C22 cells. **(C-D)** FMRP immunostaining in C22 cells **(C)** and in C22 cells treated with Fmr1 siRNA **(D)**. **(E-J)** Susceptibility of C22 cells to Nap (control (scrambled siRNA-treated, Sc) and Fmr1 siRNA-treated (Si)). **(E)** Schematic showing regimen for Nap injury. **(F-I)** Expression of markers of oxidative (4HNE) and genotoxic (γ-H2AX) stress in Sc and Si cells prior to and post Nap. **(F i-F vi)** 4HNE immunostaining (green) in Sc and Si cells prior to and post Nap. **(G)** Quantitation of 4HNE immunofluorescence per cell in Sc and Si cells prior to and post Nap (≥ 25 cells were analysed per timepoint/per experiment, n=3 experiments). **(H i-H vi)** γ-H2AX immunostaining (red) in Sc and Si cells prior to and post Nap. **(I)** Quantitation of γ-H2AX immunofluorescence per cell in Sc and Si cells post Nap. **(J)** Cytotoxicity of Nap in Sc and Si cells 24 h post Nap (n=3 experiments). Black bars (Sc), Grey bars (Si). Unpaired two-tailed t-test (p< .05*, p< .01**,p< .001***). For normality test and two-way ANOVA see Table S2. Scale Bar=5 um.

Next, we optimized methods for challenging C22 cells with Nap and for the knockdown of FMRP expression in these cells via RNA interference. Careful titration of Nap dosage and time of exposure (see methods) showed that a 1 h pulse of Nap was sufficient to induce expression of oxidative and genotoxic stress markers in C22 cells and marginally increase cell death 24 h post exposure. In an independent set of experiments, we established that treatment with 3 different Fmr1 siRNAs was sufficient to reduce FMRP levels expression by 80% or greater (see methods, compare FMRP expression in scrambled siRNA-treated cells, Sc, and Fmr1 siRNA-treated cells, Si in Fig. 2D, n>3 experiments).

To determine if FMRP regulates susceptibility to Nap in C22 cells, we incubated control (scrambled siRNA-treated cells, Sc) and FMRP-depleted (Fmr1 siRNA-treated cells, Si) cells with Nap for 1 h and the harvested cells at different timepoints for analysis (shown schematically in Fig. 2E, see methods). To assess levels of oxidative and genotoxic stress, we stained cells with anti-4HNE and anti-γ-H2AX respectively (Fig. 2F-2H). To assess Nap cytotoxicity, cells were subject to a WST-1 assay 24 h post exposure. We found that levels of 4HNE and γ-H2AX (Fig.2G-2I) were elevated in Fmr1-depleted cells at all timepoints (n=3 experiments each) and that Fmr1-depleted cells exhibited greater cell death in response to Nap (Fig. 2J). These data correlated well with the increased susceptibility of CCs to Nap in Fmr1 KO animals and demonstrated that FMRP has a cell intrinsic role in protecting cells from Nap.

### FMRP is required for the induction of the Integrated Stress Response pathway that protects from Naphthalene induced stress

The increased susceptibility of FMRP-deficient CCs and C22 cells to Nap led us to investigate further the role of FMRP in the Nap induced stress response. As mentioned previously, FMRP has been shown to regulate the formation of SGs in response to stress. SG biogenesis is an integral aspect of the stress response. However, it is currently unclear whether a defect in SG biogenesis alone can render cells more susceptible to stressful stimuli (Adjibade P et al., 2017). Nevertheless, in an effort to characterize the contribution of FMRP in the stress response, we probed SG biogenesis in control and FMRP-depleted C22 cells post Nap. TIA1, Atx2 and G3BP are integral components of SGs in mammalian cells (Anderson P & Kedersha N, 2006; Anderson P & Kedersha N, 2008). To determine whether the genesis of SGs was inhibited in FMRP-deficient C22 cells 1 h post Nap, we stained Sc and Si cells with antisera against TIA1/Atx2 (anti-TIA1 immunostaining shown in Fig. 3A, anti-Atx2 immunostaining shown in Fig. S2A). Confocal analysis showed that Sc cells contain a few sporadic SGs and the numbers of SGs increased dramatically 3, 6, and 12 h post Nap and returned to baseline by 24 h (Fig. 3A, Fig. S2A, n=3 experiments). Analysis of Si cells showed that these cells contain a few SGs in the untreated condition and that the numbers of SGs did not increase post Nap (Fig. 3A). Based on this analysis, we inferred that SG biogenesis is perturbed in FMRP-deficient cells post Nap. Co-staining cells with markers of SG and FMRP showed that FMRP has a punctate distribution both before and after Nap and that FMRP punctae were largely distinct from SGs. Based on these findings we concluded that FMRP regulates SG biogenesis in response to Nap.

**Figure 3:**
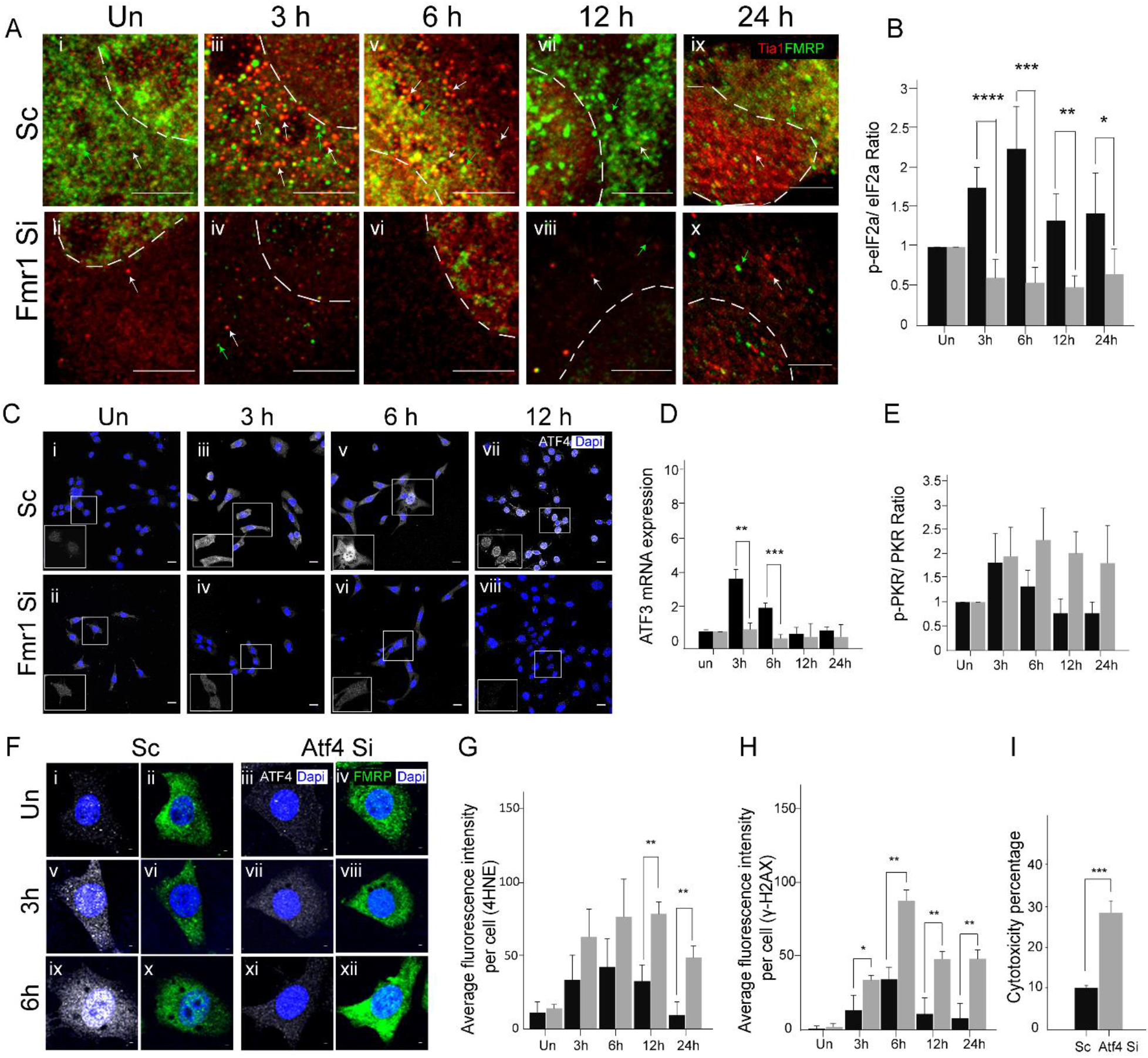
FMRP deficient C22 cells fail to upregulate the Integrated Stress Response and induce ATF4, essential for protection from Nap-induced stress. **(A i - A x)** Expression of the stress granule marker Tia-1(white, arrows) in control (Scrambled siRNA-treated, Sc) and Fmr1 siRNA-treated (Si) cells prior to and post Nap. Note that FMRP expression in the same cells (green, arrow) does not completely overlap with Tia-1. Also see Supplementary Figure 2 for Atx2/FMRP staining. **(B)** Western blot-based quantitation of phospho-eIF2α /eIF2α ratios in Sc and Si cells prior to and post Nap treatment (n=5 experiments). See Fig. S2 for representative blots used for quantitation. **(C i - C viii)** ATF4 immunostaining in Sc (upper panel) and Si cells (lower panel) prior to and post Nap. Note nuclear accumulation of ATF4 in Sc cells by 6 h post Nap (inset). See Fig. S2 for quantitation. **(D)** Quantitative RT-PCR-based analysis of expression of the ATF4 target gene ATF3 in Sc and Si cells prior to and post Nap (n=3 experiments). **(E)** Western blot-based quantitation of phospho-PKR/ PKR ratios in Sc and Si cells prior to and post Nap treatment (n=3 experiments). See Fig S2 for representative blots used for quantitation. **(F-I)** Susceptibility of C22 cells to Nap in control (Scrambled siRNA-treated, Sc) and Atf4 siRNA-treated (Si) cells. **(F i - F xii)** Analysis of ATF4 levels (white) in Sc and Si cells prior to and post Nap treatment. Immunostaining for ATF4 (white) and FMRP (green) in Sc (left panel) and Si (right panel) cells. **(G)** Quantitation of 4HNE immunofluorescence per cell in Sc and Si cells prior to and post Nap. See Fig S2 for representative images **(H)** Quantitation of γ-H2AX immunofluorescence per cell in Sc and Si. See Fig. S2 for representative images. **(I)** Cytotoxicity of Nap in Sc and Si cells 24 h post Nap exposure (n=3 experiments). Black bars (Sc), Grey bars (Si). Unpaired two-tailed t-test (p< .05*, p< .01**, p< .001***). For normality test and two-way ANOVA see Table S3. Scale Bar=5 um.

In addition to a role in SG biogenesis, FMRP also has a chromatin-dependent role in resolving certain types of genotoxic stress. More specifically, FMRP-deficient cells fail to recruit γ-H2AX to stalled replication forks and single strand breaks in response to Aphidicolin, 5HU and UV exposure but are able to recruit γ-H2AX in response to gamma-radiation (Alpatov R et al., 2014). While it is plausible that FMRP serves a similar role in Nap-treated cells, we noted that the nuclear accumulation of γ-H2AX in FMR-deficient CCs and C22 cells post Nap was greater than in the respective controls. This suggested to us that the DNA Damage Response was at least partially active in FMR-deficient cells and, more importantly, that extent of DNA damage (as reported by nuclear γ-H2AX accumulation) was greater in FMR-deficient cells than in controls (see Discussion).Together, the data led us to hypothesize FMRP has a more general role in facilitating stress responses post Nap and led to investigate its role in the Integrated Stress Response (ISR) pathway.

As previously mentioned, the ISR is induced when one of four stress-responsive kinases (GCN2, PERK, HRI, PKR) phosphorylate eIF2α at Serine 51. Phosphorylation of eIF2α arrests conventional translation, stimulates genesis of SGs and enables specialized translation of stress response proteins like ATF4 (Pakos-Zebrucka K et al., 2016; van ’t Wout EF et al., 2014). To probe the status of the ISR in C22 cells post Nap, we examined the phosphorylation state of eIF2α. We exposed C22 cells to Nap for 1 h, harvested cells at various timepoints and quantified the levels of expression of both eIF2α and phosphorylated eIF2α (p-eIF2α). Western blot-based ratiometric quantitation of total and p-eIF2α in Sc cells showed that p-eIF2α levels increased 3 and 6 h post injury and decreased to baseline levels thereafter (Fig. 3B, Fig. S2B - S2C, n=5 experiments). We inferred that the ISR is induced in C22 in response to Nap. We then exposed Si cells to Nap for 1 h and found that, contrary to controls, the levels of p-eIF2α did not increase post Nap (Fig. 3B, Fig. S2B - S2C, n=5 experiments). The analysis of p-eIF2α suggested the FMRP-depletion might inhibit the ISR.

Next, we examined the expression of ATF4 and its target, ATF3, in control and FMRP-deficient cells. Sc and Si cells were stained with an anti-ATF4 antibody prior to and post Nap. In Sc, the expression of ATF4 was undetectable in untreated cells, increased dramatically 3, 6, 12 h post Nap treatment and then approached baseline levels at 24 h (Fig. 3C, Fig. S2D, n=3 experiments). In Si cells, levels of ATF4 were negligible in untreated cells and showed no appreciable increase post Nap. Next, we assayed ATF3 levels by quantitative real-time PCR (qPCR). For this, RNA was isolated from Sc and Si cells at different timepoints and subjected it to qPCR analysis. In Sc, levels of ATF3 mRNA increased at 3, 6 h post Nap and returned to baseline thereafter (Fig. 3D, n=3 experiments). In Si, the levels ATF3 did not rise appreciably above baseline post Nap (Fig. 3D). These findings were also validated with anti-ATF3 immunostaining (data not shown). Based on these data we concluded that both ATF4 and ATF3 expression are perturbed in FMRP-deficient cells post Nap. Taken together, the findings showed that the ISR is perturbed in FMRP-deficient cells post Nap.

Next, we decided to investigate whether the upstream kinases that phosphorylate eIF2α and induce the ISR are activated (phosphorylated) in FMRP-deficient cells. We probed the expression of GCN2, PERK, HRI, PKR and their phosphorylated isoforms in Nap-treated C22 cells using commercially available antibodies (see methods). Among all pairs of antisera tested, antisera for PKR and p-PKR provided reproducible results. Western blot-based ratiometric quantitation of p-PKR and total PKR in Sc and Si cells showed that the p-PKR levels increase in both Sc and SI 3 h post Nap (Fig. 3E, Fig. S2E - S2F, n=3 experiments). Importantly, we noted that levels of p-PKR returned to baseline in Sc at 6 h and later timepoints, but remained significantly higher in Si at later timepoints (Fig. 3E). This suggested that at least one of the stress responsive kinases (PKR) is activated in FMRP-deficient cells post Nap but is unable to actuate downstream processes.

Perturbations in the ISR in FMRP-deficient C22 cells post Nap provided a plausible explanation for why these cells are more susceptible to Nap. To test this, we decided to probe how perturbing the ISR, by knocking down levels of Atf4, would impact susceptibility to Nap. Control (Scrambled siRNA) and Atf4 siRNA treated C22 cells were exposed to Nap as described earlier and cells were harvested at different timepoints for analysis. ATF4 immunostaining of control and Atf4 siRNA-treated cells showed that siRNA treatment eliminated ATF4 expression in cells post Nap exposure (Fig. 3F). We also found that ATF4-depleted cells exhibited increased expression of 4HNE (Fig. 3G, representative images shown in Fig. S2G) and γ-H2AX (Fig. 3H, representation images shown in Fig. S2H) at all timepoints examined (n=3 experiments each, quantitation of cell fluorescence based on n>25 cells per experiment) and also increased cell death 24 h post injury (Fig. 3I). We concluded that the increased levels of oxidative and genotoxic stress and increased cytotoxicity observed in FMRP-deficient cells could be due to a failure to induce the ISR and upregulate ATF4.

In light of the findings in C22 cells, we examined whether perturbations to the ISR are also observed in FMRP-deficient CCs in Nap-treated mice. We counterstained sections from control and Fmr1 KO lungs post Nap with antisera to both ATF4 and ATF3. Although ATF4 immunostaining was inconclusive, we noted that the levels of ATF3 were negligible in CCs in the control lung and upregulated post Nap (Fig. S2I - S2J, sections from n=3 mice). Pertinently, the levels of ATF3 in CCs in Fmr1 KO did not increase post Nap. These results are consistent with a role for FMRP in the induction of ISR in CCs post Nap.

### FMRP is expressed in the airways of the human lung and protects human bronchial BEAS-2B cells from 9, 10-Phenanthrenequinone induced stress

The findings in the murine lung led us to ask whether FMRP has a conserved role in the human lung. To investigate this possibility, we first examined the distribution of FMRP in the human lung. Paraffin sections stained with FMRP antisera showed that FMRP is expressed throughout the airways and more broadly (Fig. 4A, n=2 sections each from n=5 independent lung biopsies). Triple labeling experiments with markers for ciliated cells and CCs showed that FMRP is expressed in both ciliated and non-ciliated cells, including CCs. Based on the distribution of FMRP we surmised that the protein could play a role in the airways in the human lung as well.

**Figure 4:**
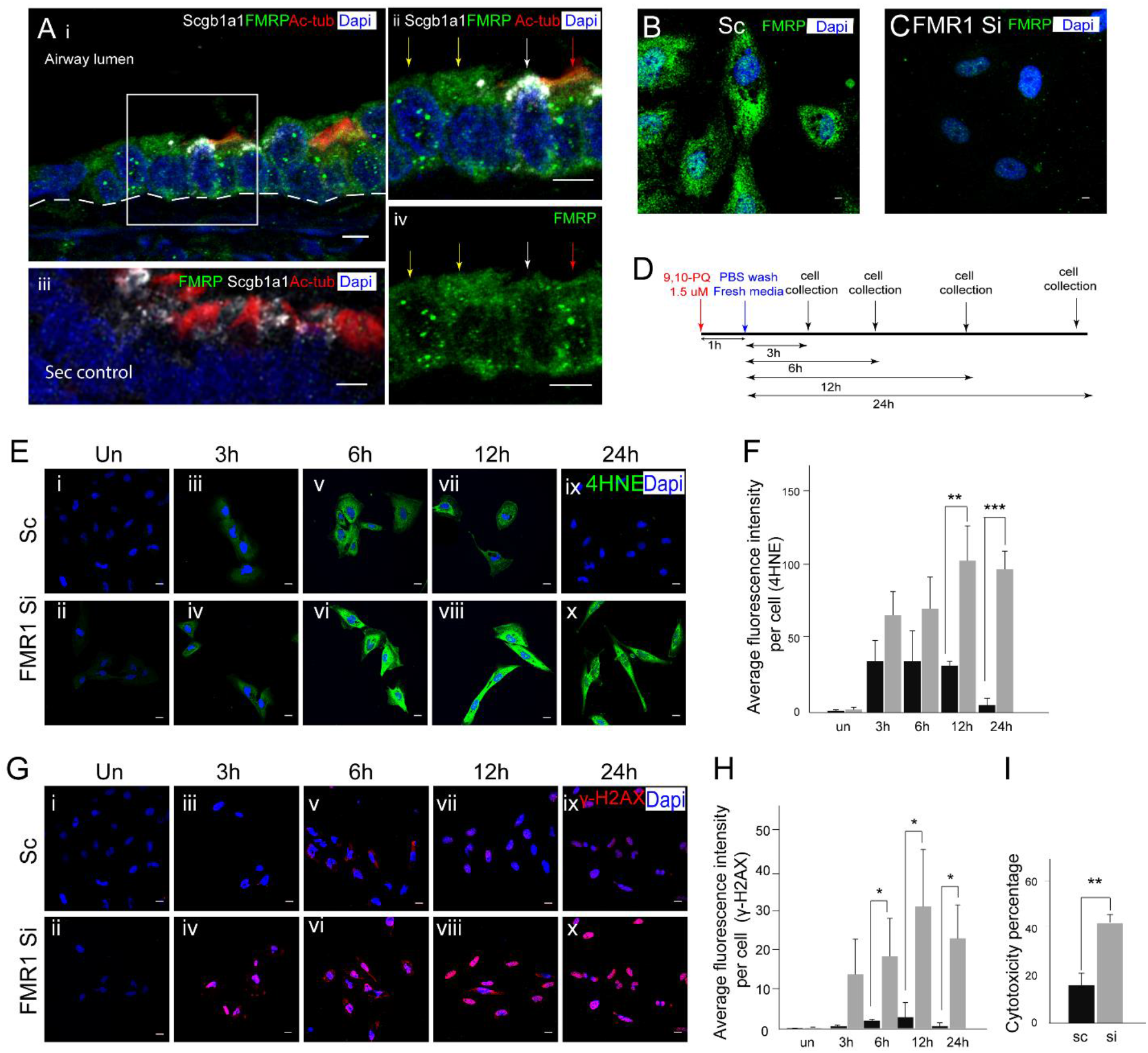
FMRP is expressed in the human airways and protects human bronchial BEAS-2B cells from 9, 10-Phenanthrenequinone-induced stress. **(A-C)** FMRP expression in the human lung and in BEAS-2B cells, a cell line derived from the human bronchial epithelium. **(A i - A iv)** FMRP immunostaining (green) in the distal airways of the human lung. **(A i, A ii, A iv)** Stained section showing FMRP expression in airway non-ciliated cells (Scgb1a1+ (white), white arrows; Scgb1a1-, yellow arrows) and ciliated cells (red, red arrow). Boxed area shown at higher magnification in top and bottom panels on the right. Negative control (secondary alone) for FMRP immunostaining shown in **A iii**. **(B)** FMRP immunostaining of BEAS-2B cells. **(C)** FMRP immunostaining of FMR1 siRNA-treated BEAS-2B cells. **(D-I)** Susceptibility of BEAS-2B cells to PQ injury in control (Scrambled siRNA-treated, Sc) and Fmr1 siRNA-treated (Si) cells. **(D)** Schematic showing regimen for PQ injury. **(E-H)** Expression of markers of oxidative (4HNE) and genotoxic (γ-H2AX) stress in Sc and Si cells prior to and post PQ. **(E i - E x)** 4HNE immunostaining (green) in Sc and Si cells prior to and post PQ. **(F)** Quantitation of 4HNE immunofluorescence per cell in Sc and Si cells post PQ (≥ 25 cells were analysed per timepoint/per experiment, n=3 experiments). **(G i - G x)** γ-H2AX immunostaining (red) in Sc and Si cells prior to and post PQ. **(H)** Quantitation of γ-H2AX immunofluorescence per cell in Sc and Si cells post PQ. **(I)** Cytotoxicity of PQ in Sc and Si cells 24 h post PQ exposure (n=3 experiments). Black bars (Sc), Grey bars (Si). Unpaired two-tailed t-test (p< .05*, p< .01**, p< .001***). For normality test and two-way ANOVA see Table S4. Scale Bar=5 um.

The BEAS-2B cell line is derived from normal human airways. These cells do not express markers of ciliated cells and, akin to CCs, have characteristics of non-ciliated cells. We stained BEAS-2B cells with FMRP antisera to find that these cells expressed FMRP (Fig. 4B, n>6 experiments). We then proceeded to develop an assay to probe the role of FMRP in stress responses in these cells.

Since the susceptibility of airway CCs to Nap is not recapitulated in the human lung or in BEAS-2B cells (data not shown), we utilized a different injury model to probe the role of FMRP in stress responses in human cells. 9, 10-Phenanthrenequinone (PQ) is an air pollutant that is present at high levels in diesel exhaust particles and is known to trigger oxidative and genotoxic stress (Lavrich KS et al., 2018). As part of our characterization of PQ, we first exposed control (scrambled siRNA-treated cells) and FMRP-depleted (Fmr1 siRNA-treated cells) C22 cells to a pulse of PQ for 1 h (see methods) and harvested cells at different timepoints for analysis (shown schematically in Fig. S3A). Consistent with our findings in the Nap model, we found that FMRP-depleted C22 cells exhibited increased expression of 4HNE (Fig. S3B) and γ-H2AX (Fig. S3C) and increased cell death 24 h post exposure (Fig. S3D). We then examined ATF4 expression to find that although ATF4 levels increased in Sc cells at 3 h, 6 h post PQ, no expression was detected in Si cells (Fig. S3E). These experiments showed that PQ treatment does lead to oxidative and genotoxic stress and that, FMRP-deficient C22 cells are more susceptible.

Next, we exposed control (scrambled siRNA-treated cells) and FMRP-depleted (FMR1 siRNA-treated cells) BEAS-2B cells to a pulse of PQ for 1 h and harvested them at different timepoints for analysis (shown schematically in Fig. 4D). We determined independently that the protocol for the knockdown of FMRP in BEAS-2B cells lead to a 90% reduction in the levels of FMRP post treatment (Fig. 4C, n>3 experiments). We found that FMRP-depleted cells exhibited increased expression of 4HNE (Fig. 4E-4F) and γ-H2AX (Fig. 4G-4H) at all timepoints examined (n=3 experiments each, quantitation of cell fluorescence based on n>25 cells per experiment) and increased cell death 24 h post injury (see methods, Fig. 4I). These experiments showed that FMRP-deficient BEAS2B cells are more susceptible to PQ.

### FMRP is required for the induction of the Integrated Stress Response pathway that protects from 9, 10-Phenanthrenequinone induced stress

Next, we determined whether FMRP is required for the induction of the ISR in BEAS-2B cells. As described previously, we probed the activation status of PKR (ratiometric quantitation of p-PKR and total PKR in both control cells (Scrambled siRNA-treated, Sc) and FMRP-deficient cells (Fmr1 SiRNA-treated, Si) Fig. 5A, Supplementary Fig. 4A-4B, n=3), the phosphorylation status of eIF2α (Fig. 5B, Fig. S4C - S4D, n=5), the formation of SGs (Fig. 5C, Fig. S4E n=3), the levels of ATF4 induction (Fig. 5D-5E, n=3) and the levels of ATF3 induction (Fig. 5F, n=3) at different times post PQ. These experiments showed that although p-PKR levels were increased in both Sc and Si post PQ, all of the downstream processes of the ISR were perturbed in Si. We concluded that FMRP is required for the ISR in BEAS-2B cells post PQ.

**Figure 5:**
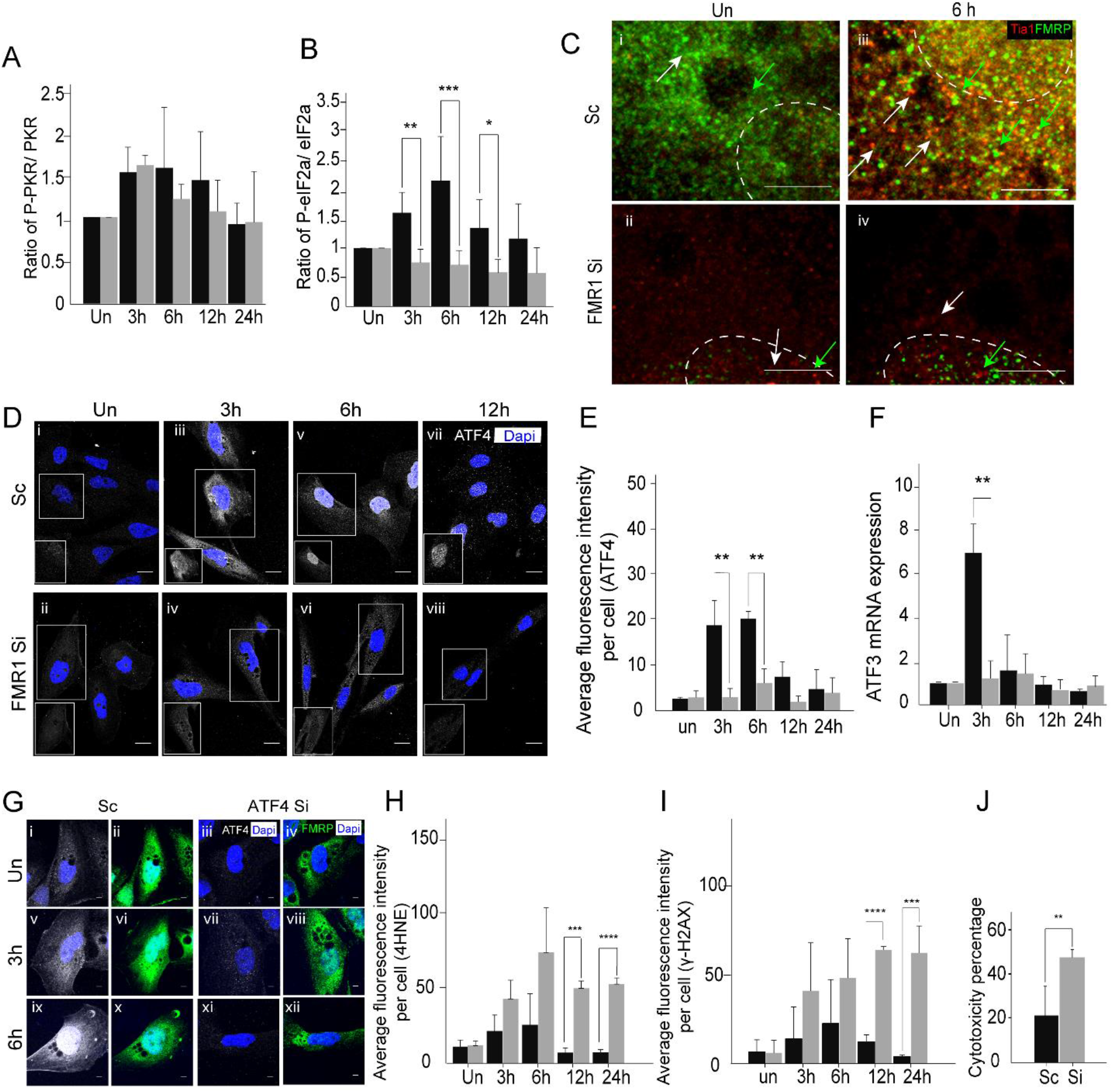
FMRP deficient BEAS-2B cells fail to upregulate the Integrated Stress Response and induce ATF4, essential for protection from 9, 10-Phenanthrenequinone induced stress. **(A)** Western blot-based quantitation of phospho-PKR/ PKR ratios in Sc and Si cells prior to and post Nap treatment (n=3 experiments). See Fig. S4 for representative blots used for quantitation. **(B)** Western blot-based quantitation of phospho-eIF2α /eIF2α ratios in Sc and Si cells prior to and post PQ treatment (n=5 experiments). See Fig. S4 for representative blots used for quantitation. **(C i-C iv)** Expression of the stress granule marker Tia-1(white, arrows) in control (Scrambled siRNA-treated, Sc) and Fmr1 siRNA-treated (Si) cells prior to and post PQ. Note that FMRP expression in the same cells (green, arrow) does not completely overlap with Tia-1. Also see Fig. S4 for G3BP/FMRP staining. **(D-E)** Analysis of ATF4 prior to and post PQ. **(D)** ATF4 immunostaining in Sc (upper panel) and Si cells (lower panel) prior to and post PQ treatment. Note nuclear accumulation of ATF4 in Sc cells by 6 h post PQ treatment (inset). **(E)** Quantitation of ATF4 immunofluorescence per cell in Sc and Si cells prior to and post PQ. **(F)** Quantitative RT-PCR-based analysis of expression of ATF3 in Sc and Si cells prior to and post PQ (n=3 experiments). **(G-J)** Susceptibility of BEAS-2B cells to PQ in control (Scrambled siRNA-treated, Sc) and Atf4 siRNA-treated (Si) cells. **(G i - G xii)** Analysis of ATF4 levels (white) and FMRP (green) in Sc (left panel) and Si (right panel) cells prior to and post PQ treatment. Immunostaining for ATF4 (white) and FMRP (green) in Sc (left panel) and Si (right panel) cells. **(H)** Quantitation of 4HNE immunofluorescence per cell in Sc and Si. See Fig. S4 for representative images. **(I)** Quantitation of γ-H2AX immunofluorescence per cell in Sc and Si. See Fig. S4 for representative images. **(J)** Cytotoxicity of PQ in Sc and Si cells 24 h post PQ treatment (n=3 experiments). Black bars (Sc), Grey bars (Si). Unpaired two-tailed t-test (p< .05*, p< .01**, p< .001***). For normality test and two-way ANOVA see Table S5. Scale Bar=5 um.

Next, we investigated whether the loss of ATF4 would recapitulate the loss of FMRP post PQ. Control (Scrambled siRNA) and ATF4 siRNA treated BEAS-2B cells were exposed to PQ as previously described and cells were harvested at different timepoints for analysis. Consistent with expectations, ATF4 siRNA-treated cells showed no anti-ATF4 immunostaining post PQ exposure (Fig. 5G, n>3 experiments). We found that ATF4-depleted BEAS-2B cells exhibited increased expression of 4HNE (Fig. 5H, representative images shown in Fig. S4F) and γ-H2AX (Fig. 5I, representative images shown in Fig. S4G) and increased cell death 24 h post injury (see methods, Fig. 5J). These data indicated that the loss of ATF4 largely phenocopies the loss of FMRP in PQ-treated BEAS-2B cells.

## DISCUSSION

The aim of this study was to probe the role of FMRP in stress responses in the lung. We report that FMRP plays an essential role in protecting the airways in mice, and potentially in humans, from the deleterious effects of xenobiotic stress. Our studies provide strong evidence that FMRP protects the lung by facilitating the induction of the ISR (see model, Fig. 6). In the paragraphs that follow we will discuss the plausible mechanism/s by which FMRP may regulate the ISR, the possibility that FMRP regulates stress response pathways in addition to the ISR, and the clinical implications of the findings reported here.

**Figure 6:**
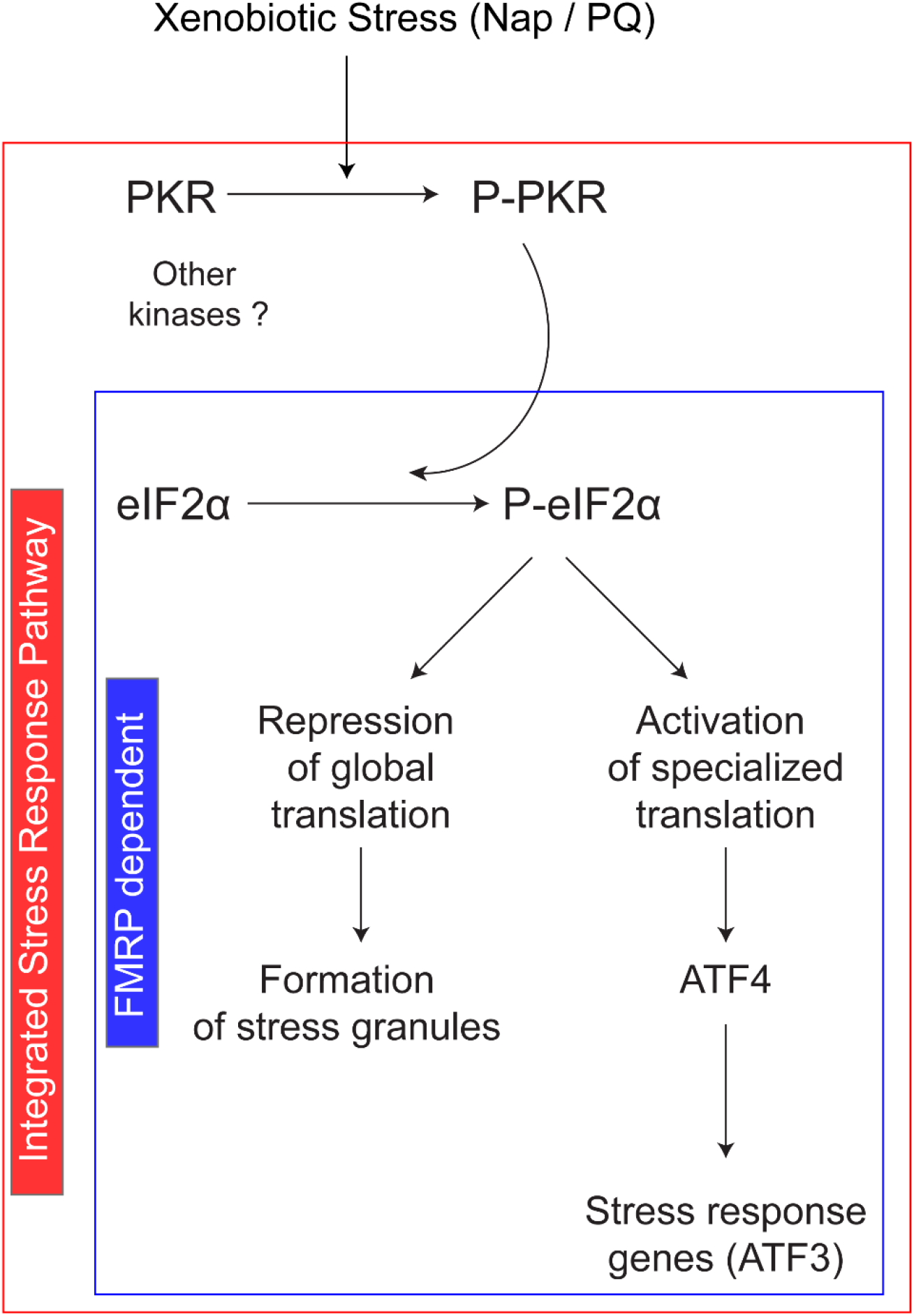
Model for the role of FMRP in the regulation of the Integrated Stress Response in the lung. Exposure to xenobiotics such as Naphthalene (Nap) and 9,10-Phenanthrenequinone (PQ) result in the activation of at least one of four stress-responsive kinases (PKR) and to the induction of Integrated Stress Response Pathway (ISR, outlined in red). Our findings suggest that FMRP has an essential role downstream to PKR phosphorylation (outline in blue).

A major finding of our study is that FMRP is required for the actuation of the ISR pathway. More specifically, we find that the stress responsive kinase PKR is activated in FMRP-deficient cells but that the phosphorylation of the PKR substrate, eIF2α, is perturbed. The mechanism by which FMRP regulates this step is currently unknown. The analysis of FMRP-binding proteins in neuronal and other tissues has identified numerous interacting partners. Interestingly, among these interacting partners are the proteins Caprin1 and G3BP that have independently been implicated in the induction of the ISR pathway in response to stress (Taha MS et al., 2020; Wu Y et al., 2016). Pertinently, both Caprin1 and G3BP1 have been shown to be important for eIF2α phosphorylation (Reineke LC et al., 2015). Thus, it is plausible that FMRP acts in concert with Caprin1 and G3BP1 to facilitate eIF2α phosphorylation. Although eIF2α phosphorylation is an early event in the ISR pathway and perturbations at this stage are likely to affect all downstream processes, our data do not allow us to rule out the possibility that FMRP has independent roles in downstream processes. FMRP could, for example, also have an independent role in stress granule biogenesis (Didiot MC et al., 2009; Linder B et al., 2008). Our future experiments will probe these possibilities.

Studies that have examined the role of FMRP vis-à-vis stress responses suggest that FMRP could protect cells from stress in myriad ways. For example, it has been demonstrated that FMRP plays a chromatin-dependent role in inducing the DDR. This could be relevant in the context of the lung. Along the same lines, there is also evidence that FMRP regulates the expression of Superoxide Dismutase1 (SOD1) in the brain (Bechara EG et al., 2009). Levels of SOD1 are reduced in the brains of Fmr1 KO animals. Since SOD1 has an important role in protecting cells from stress, FMRP could alter the susceptibility of tissues to stressful stimuli by altering the baseline levels of SOD1. To investigate this possibility, we probed levels of SOD1 in the brain and lung using both Western Blot and immunohistochemical approaches (Fig. S5A). Although we did observe that SOD1 levels in the brain were lower in Fmr1 KO than wild type, the levels of SOD1 in the lung were comparable. Moreover, we also analyzed SOD1 levels in the bronchial cell lines (C22 and BEAS-2B) with or without FMRP to find that SOD1 levels were comparable (Fig. S5D - S5I). Taken together, these data show FMRP is unlikely to regulate SOD1 expression in the lung. Nevertheless, the role of FMRP in the DDR, and in the regulation of SOD1 expression, show that FMRP can contribute toward protecting tissues from stress by ISR-independent mechanisms as well.

Although FMRP is broadly expressed in humans and mice alike (https://www.genecards.org/cgi-bin/carddisp.pl?gene=FMR1), historically, FMRP has almost exclusively been studied in a neural context due to its connection with intellectual disability. An important finding of this study is that it demonstrates a role for FMRP in the lung and points to a potential vulnerability in individuals with an FMR1 deficiency. Clinically, the bulk of the case studies on FXS patients are derived from geographic regions where the load of pulmonary environmental stressors is low. Our study suggests that individuals with FXS living in areas of higher pollutant load may be more susceptible to lung damage/disease and FMRP status in the lung may be a strong correlate of resilience to pulmonary insults.

## MATERIALS AND METHODS

All animal work reported here has been approved by the Internal Animal Users Committee (IAUC) and the Institutional Animal Ethics Committee (IAEC) at inStem. Any procedure that could conceivably cause distress to the animals employed pre-procedural anesthesia with isofluorane gas (Baxter Healthcare Corp.), delivered by an anesthetic vaporizing machine. All animals were monitored for signs of distress and euthanized if in distress. The analysis of human biopsies was approved by Institutional Ethics Committee of JSS Medical College.

### Mouse strains

Fmr1 knockout mice (*Mus musculus*) strain was maintained on a C57B6/J background at the Centre for Brain Development and Repair (CBDR), inStem. Genotyping of the animals was done using established protocols (Bakker CE et al., 1994).

### Human samples

Human (*Homo sapiens*) lung tissue was obtained from five subjects at autopsy by a forensic pathologist from JSS Medical College, Mysore. The cause of death was not attributed to lung trauma. Deidentified samples were fixed in 4% paraformaldehyde at 4 °C overnight, embedded in paraffin, and processed for immunohistochemical analysis.

### Cell lines and culture conditions

Human lung (BEAS-2B) non-ciliated airway epithelial origin cell line was obtained from Johns Hopkins University (kind gift from Prof. S. Biswal) (Singh A et al., 2009). The murine Club cell line (C22) was purchased from ECACC, UK (cat no.07021401, #07D022). Both cell lines were tested for mycoplasma contamination and found to be negative. BEAS-2B were grown in DMEM: F12K (Gibco, USA, 21127030) (1:1) media, supplemented with 10% FBS (Gibco,USA, 10082147) and Penicillin-Streptomycin (Gibco, USA, 1540122) at 37°C, 5% CO2. C22 cell line was maintained in a proliferative state as per suppliers’ instructions and experiments were performed 24 h post differentiation. Experiments were conducted within 3rd to 7th passages for BEAS-2B and within 3rd to 12th passages for C22.

### Models for xenobiotic stress

For Naphthalene (Nap) injury in mice, wild type or Fmr1 knockout mice aged (≥8 weeks of age) were injected intraperitoneally with Corn Oil (vehicle, Sigma, USA, C8267) or with Nap dissolved in corn oil (300 mg kg^−1^, (Sigma, USA, 147141) using established protocols (Guha A et al., 2014; Guha A et al., 2012). Animals were sacrificed 12 h, 24 h, 48 h after injection for analysis.

To establish an assay for Nap injury in C22 cells, we first determined that these cells expressed the cytochrome Cyp2f2 that converts Nap to stress-inducing derivatives. Having established this, we tested a range of concentrations of Nap (50 ug ml^−1^ to 500 ug ml^−1^, in DMSO/DMEM). Nap was found to be stable in solution at concentrations upto 100 ug mL^−1^ and unstable at higher concentrations leading to cell death within 3h post exposure. Nap exposure at 50-75 ug ml^−1^ (DMSO/DMEM, DMSO final concentration 0.7%) for short (1 h) and long duration (24 h) led to a progressive increase in expression of stress markers and mild cytotoxicity after a 24 h period. To probe the effects of FMRP/ATF4 deficiency on susceptibility to Nap, cells were exposed to Nap at 75 ug ml^−1^ (DMSO/DMEM, DMSO final concentration 0.7%) for a period 1h. Cells were then washed in PBS and chased for varying periods of time in complete medium.

It has been reported previously that 9,10-Phenanthrenequinone (PQ) causes a sharp decrease in the viability of BEAS-2B cells when administered to cells for 24 h at concentrations greater than 1 uM (Koike E et al., 2014). We reconfirmed these findings and determined the LD50 dose to be ~1.5 uM ((Sigma, USA, 275034), dissolved in DMSO/DMEM, DMSO final concentration 0.00002%). To probe the effects of FMRP/ATF4 deficiency on susceptibility to PQ, cells were exposed to PQ at 1.5 uM (DMSO/DMEM, DMSO final concentration 0.00002%) for a period 1h. Cells were then washed with PBS and fresh complete media and chased for varying period’s time in complete medium.

### siRNA based knockdown of FMR1/ATF4 expression

Several studies have demonstrated that multiple siRNA administered together or sequentially work more efficiently for silencing gene expression than a single siRNA (Wang Z et al., 2016; Fähling M et al., 2009; Zhang P et al., 20015; Hatch EM et al., 2010). For our studies we used 3 distinct siRNAs for each targeted gene. siRNAs were administered to cells sequentially, 12 h apart, to silence the gene expression. siRNA transfections were done with Lipofectamine 2000 (Thermofisher Scientific, USA, 11668027). All xenobiotic stress assays in C22 cells were performed 36 h after treatment with the last siRNA. C22 cells were transferred from proliferative to differentiation-inducing media 12 h after the last Si RNA treatment and utilized for xenobiotic stress assays 24 h thereafter. All xenobiotic stress assays in BEAS-2B cells were performed 12 h after treatment with the last siRNA. All siRNAs were obtained from Ambion: murine Fmr1 (Ambion, USA, 4390771), murine Atf4 (Ambion, USA, 16708), human FMR1 (Ambion, USA, 4392420) and human ATF4 (Ambion, USA, 16708), and Scrambled (Negative control, Ambion, USA, 4390843). The assay IDs for each of siRNAs are as follows: Mouse Fmr1 siRNA (Assay ID: 5315, 5317, s66177), Human FMR1 siRNA (Assay ID: 5315, 5316, 5317), Mouse Atf4 siRNA (Assay ID: 160775, 160776, 160777), Human ATF4 siRNA (Assay ID: 122168, 122287, 122372).

### Cell Cytotoxicity assay

C22 and BEAS-2B cells were inoculated into 96-well plate and treated with Nap or PQ for 1h, as described above, and harvested for analysis 24 h later. Cell viability was assayed using WST-1 reagent (Sigma, USA, 5015944001)). Briefly, cells were incubated with WST-1 for 4h and absorbance readings were taken and analyzed as per manufacturer’s protocols. Cytotoxicity percentage=100 X [(OD (450nm-650nm) of untreated cells-OD (450nm-650nm) of treated cells)/ OD (450nm-650nm) of untreated cells].

### Histology, Immunofluorescence and Imaging

Lungs were inflated with 4% (wt/vol) Paraformaldehyde (Alfa Aesar, USA, 30525-89-4) in PBS and fixed for 8 hours at 4°C. Fixed lungs were subsequently embedded in paraffin, sectioned (5 um) and processed for immunohistochemical analysis post heat-mediated antigen retrieval at pH 6.0 (Vector Labs, USA) except sections stained with anti-SOD1 antisera that were subject to antigen retrieval at pH 9.0 (Vector Labs, USA). For cellular immunostaining, cells were seeded on coated coverslips (0.1% gelatin, Sigma, USA, G9391, as per manufacturer’s protocol). Post-treatment, cells were fixed with 4% PFA for 30 minutes and blocked with 2%FBS, 0.2% BSA and 0.1% Triton X 100 in 1X PBS for an hour and stained. Primary antibodies were diluted using the same blocking solution. Immunohistochemical analysis utilized the following antisera: rabbit anti-FMRP (Abcam, UK, 17722, 1:500), rabbit anti-FMR1 (Sigma, USA, 1:200), goat anti-Scgb1a1 (Santa Cruz, USA, Sc365992, 1:500), mouse anti-acetylated tubulin (Sigma, USA, T7451, 1:1000), mouse anti-4HNE (Abcam, UK, ab48506, 1:500), rabbit anti- γ-H2AX (Novus biological, USA, NB100-384, 1:1000), mouse anti-Cyp2f2 (Santa Cruz, USA, 1:200), mouse anti-ATF4 (Sigma, USA, WH0000468M1, 1:200), goat-Anti-Tia1 (Santa Cruz, USA, SC1751, 1:200), and Alexa 488/568/647-conjugated donkey anti-mouse/rabbit/goat secondary antibodies (Invitrogen, USA, 1:300). Stained sections were mounted in ProLong Diamond (Invitrogen, USA, P36962). All samples were imaged on a FV3000 4-laser confocal microscope or on a Zeiss LSM-780 (Carl Zeiss AG, Germany) laser-scanning confocal microscope.

### Quantitative fluorescence microscopy

Frequencies of Club cells/mm airway, and total cellular fluorescence in Club cells, in lung sections, were determined from single tiled optical sections acquired on a confocal microscope using ImageJ software. For Club cell frequency analysis, cells attached to the basement membrane were counted per section per animal. Total cellular fluorescence intensity was calculated by subtracting a “background” value per section from the integrated density per cell (outlined using the software) (for FMRP, 4HNE, γ-H2AX and ATF4). The “background” value was determined by sampling integrated density of regions on the section devoid of cells. Total cellular fluorescence of C22 and BEAS-2B cells was estimated from single optical sections on a confocal microscope using ImageJ software. In all experiments involving C22 and BEAS-2B cells, ≥25 cells were analyzed per timepoint, per experiment, n=3 experiments. The images of Scgb1a1 and Cyp2f2 expression in C22 cells, and of stress granule markers in BEAS-2B and C22 cells, are maximum intensity projection images of z-stacks acquired on confocal microscope.

### Western blot analysis

Protein was extracted from cell lysates using RIPA buffer (Thermofisher Scientific, USA, 89900) containing Sigmafast EDTA free protease inhibitor cocktail (Sigma, USA, s8830) and Phosstop (Merk, USA, 4906845001). Total protein was run on a 12% SDS PAGE, transferred onto a nitrocellulose membrane (Amersham, UK, 10600002), and the membrane was stained with reversible MemCode (Thermofisher Scientific, USA, 24580) for total protein estimation (imaged on ImageQuant600 (Amersham, UK) and quantified using ImageJ). The membrane was subsequently de-stained, blocked with 5% BSA (Sigma, USA, A9418) for 1 h and probed using the following primary antisera: rabbit anti-Phospho-eIF2α (Ser51) (Cell Signalling Technology, USA, 9721S, 1:1000), mouse anti-eIF2α (Cell Signalling Technology, USA, 2103S, 1:1000), rabbit anti-Phospho-PKR (Sigma, USA, SAB4504517,1:3000), mouse anti-PKR (Santa Cruz, USA, Sc-6282, 1:1000), rabbit anti-GCN2 (Cell Signalling Technology, USA, 3302s), mouse anti-Phospho-GCN2 (Cell Signalling Technology, USA, 3301S), rabbit anti-PERK (Cell Signalling Technology, USA, 3192s, 1:1000), rabbit anti-Phospho-PERK (Cell Signalling Technology, USA, 1379s), mouse anti-HRI (Santa Cruz, USA, sc-365239). Primary antisera was detected using the following secondary antisera: HRP-conjugated anti-rabbit (abcam, UK, 6721, 1:3000) and HRP-conjugated anti-mouse secondary (Invitrogen, USA, # 62-6520 1:5000) antibodies and ECL (BioRad, USA, 1620177) and analyzed (imaged on ImageQuant600 (Amersham, UK) and quantified using ImageJ). The levels of eIF2α, phospho-eIF2α, PKR and Phospho-PKR were normalized to the total protein content of the respective lanes.

### Quantitative PCR (qPCR) analysis

RNA from cell lysates was extracted using Trizol (Invitrogen, USA, 15596018, as per manufacturer’s protocol) and qPCR was performed using the primers listed in the following table. The qPCR assays were constituted with the Maxima SYBR green/ROX qPCR Mastermix (2X) (Thermo Scientific, USA, K0221) and analyzed on a BioRad CFX3 real-time PCR system (BioRad, USA).

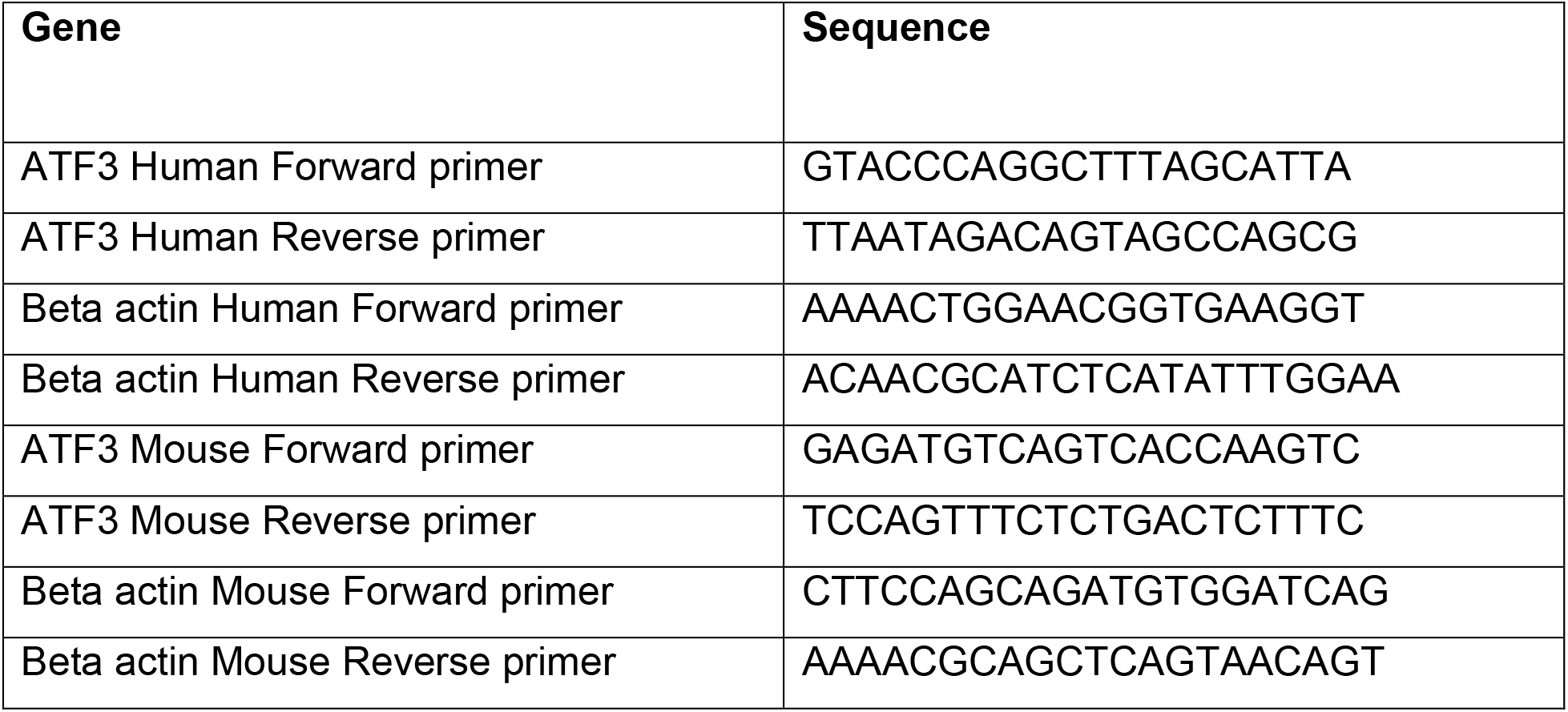

### Statistical Analysis

Statistical significance of datasets was assessed using unpaired two-tailed t-tests post Shapiro-Wilk tests for normality. Data were also analysed using a two-way ANOVA to compare changes in two groups with respect to time, genotype and interaction parameters. ANOVA data and normality test results for each figure are presented (Table S1 - S9) in a tabular format.

## ACKNOWLEDGEMENTS

We thank Joseph Jomon, National Centre for Cell Science (NCCS), Pune for sharing antibodies; Aditya Deshpande, inStem for assistance in animal experiments; Sarfaraz Nawaz and Sudhriti Ghosh Dastidar, inStem for their assistance with biochemical analyses; Harlin Kaur, Binita Dam, Arnab Karmakar, Saraswati Chavda and Mamta Yadav for technical assistance; and the Central Imaging and Flow Cytometry Facility (CIFF) and Animal Care and Resource Center (ACRC) Facility at Bangalore Life Science Cluster (BLiSC) for their constant support.

